# Neurochemical markers of uncertainty processing in humans

**DOI:** 10.1101/2025.02.19.639013

**Authors:** Nazia Jassim, Peter Thestrup Waade, Owen Parsons, Frederike H Petzschner, Caterina Rua, Christopher T Rodgers, Simon Baron-Cohen, John Suckling, Christoph Mathys, Rebecca P Lawson

## Abstract

How individuals process and respond to uncertainty has important implications for cognition and mental health. Here we use computational phenotyping to examine individualised “uncertainty fingerprints” in relation to neurometabolites and trait anxiety in humans. We introduce a novel categorical state-transition extension of the Hierarchical Gaussian Filter (HGF) to capture implicit learning in a four-choice probabilistic sensorimotor reversal learning task by tracking beliefs about stimulus transitions. Using 7-Tesla Magnetic Resonance Spectroscopy, we measured baseline neurotransmitter levels in the primary motor cortex (M1). Model-based results revealed dynamic belief updating in response to environmental changes. We further found region-specific relationships between M1 glutamate+ glutamine levels and prediction errors and volatility beliefs, revealing an important neural marker of probabilistic reversal learning in humans. High trait anxiety was associated with faster post-reversal responses. By integrating computational modelling with neurochemical assessments, this study provides novel insights into the neurocomputations that drive individual differences in processing uncertainty.

## Introduction

Navigating uncertain environments presents a unique challenge to the human brain, and by extension, mental health. The ability to extract statistical regularities from the sensory environment is of paramount importance to cognition and overall development ^1–4^. This is considerably more challenging in unpredictable or volatile environments. Here we are required to constantly adapt to the changing statistics and to update our expectations based on unexpected outcomes ^5–7^. To deal with this challenge, we estimate different types of uncertainty and use these estimates to guide our actions ^8–10^. When faced with similar contexts, individuals may process and respond to uncertainty differently, suggesting differences in their beliefs or estimates of how uncertain the environment is. Altered learning under uncertainty has been found to predict mental health difficulties, especially anxiety ^11–20^. Consequently, it is crucial to not only understand the neurocomputations of uncertainty processing, but also to use computational phenotyping ^21–24^ to identify hidden variables that account for individual differences — what we refer to as “uncertainty fingerprints.”

While there has been significant progress in the computational modelling of adaptive learning and choice behaviours, experimental paradigms and models have mainly been limited to binary choices and outcomes. However, in the real world, learning and decision-making under uncertainty are far more complex. In their seminal work on the multi-armed bandit task, Daw et al. (2006) showed that presenting individuals with four options – as opposed to two – significantly increased the levels of uncertainty, prompting more complex strategies for tracking and adapting to changing reward contingencies^25^. In a similar vein, but moving beyond reward-based learning, we examine how individuals implicitly learn the probabilistic structure of a four-choice sensorimotor learning task and how they respond to volatility as induced by a reversal of probabilities. While studies have found that volatility *increases* learning ^5,10,26–28^, the complex interplay between experimentally induced uncertainty and non-binary choices is unclear. To account for this, we present a novel instantiation of a hierarchical Bayesian model for adaptive learning that captures how people implicitly build and update beliefs about transition probabilities across four possible options.

In simplest terms, learning is driven by prediction errors - the difference between what was expected and what actually occurred ^29,30^. Minimising these prediction errors – as is central to predictive coding models ^31^ - may be equivalent to performing Bayesian inference ^32,33^. Here unsigned prediction errors can be quantified as the "surprise" (ℑ) - the negative (log) probability of an observation occurring given expectations ^9,34^. The Hierarchical Gaussian Filter (HGF), a Bayesian predictive coding model for adaptive learning in uncertain environments, was designed to match the human brain’s belief-updating process ^6,35^. HGF models using continuous and binary observations and responses have been widely used in neuroscience and psychology. The HGF estimates hidden states of the environment based on noisy observations; these estimates are analogous to the agent’s “beliefs”. It operates hierarchically in that each level models beliefs about the level below, with the lowest level representing the raw sensory data and higher levels capturing more abstract beliefs. The model is generative; specifically, it is a model of how the environment generates the observations received by the agent, with each layer in the model generating sufficient statistics of the level below. The original instantiation of the HGF has been extended to a network formulation that includes value coupling between hierarchical belief states ^36^. Here, beliefs at different levels are represented by probabilistic nodes, with higher-level nodes predicting the value (i.e, the drift) and volatility (i.e rate of change) of lower-level nodes, in what is referred to as value- and volatility-coupling respectively.

The brain uses probabilistic models to optimise performance and guide responses during sensorimotor learning ^31,33,37^. This is reflected in the motor domain as faster or more accurate responses. As sensory evidence is accumulated, it is prominently represented in motor cortical areas to guide task-relevant actions ^38,39^. At the cellular level, as learning progresses, primary motor cortex neurons (M1) neurons undergo adaption and glutamate-driven plasticity ^40,41^. Specifically, the primary excitatory neurotransmitter glutamate plays a critical role in rapid inter-cellular communication to facilitate learning and choice behaviour ^42^. Research in rodents has shown that adaptive learning in dynamic environments—as measured through reversal learning paradigms—is strongly dependent on glutamatergic modulation ^43–46^. With the advent of Magnetic Resonance Spectroscopy (MRS), a non-invasive technique to measure tissue metabolites *in vivo,* the metabolites related to learning can now be studied effectively in living humans ^47,48^. How uncertainty is encoded in the brain is an ongoing area of investigation and more work is needed to understands its neural substrates ^7,8,49,50^.

In this study, we use a combination of experimental psychology, computational cognitive modelling, and brain imaging to examine “uncertainty fingerprints” in relation to neurochemistry in humans. We capitalise on the heightened sensitivity of 7-Tesla (7T) MRS to measure excitatory glutamate + glutamine (Glx) and inhibitory *γ*-aminobutyric acid (GABA) in the primary motor cortex (M1). We implement a four-choice probabilistic sensorimotor learning task with a reversal to gauge how individuals implicitly learn the underlying statistics of the task, and how this learning is updated following a switch to the probabilities (*Fig 1*). As the learning is implicit, it can be gauged through motor responses. We further probe the latent variables and hidden states underpinning task performance by means of computational modelling. Notably, we present the first-ever implementation of a categorical state-transition HGF to model probabilistic learning – as assessed by reaction times – in the well-established four-choice Serial Reaction Time (SRT) task ^3^ (*Fig 1*). We further examine the relationships between computational model-based parameter estimates of sensorimotor probabilistic reversal learning and neurotransmitter levels. Finally, we test the hypothesis that beliefs about volatility are dependent upon trait anxiety levels.

**Figure 1.**
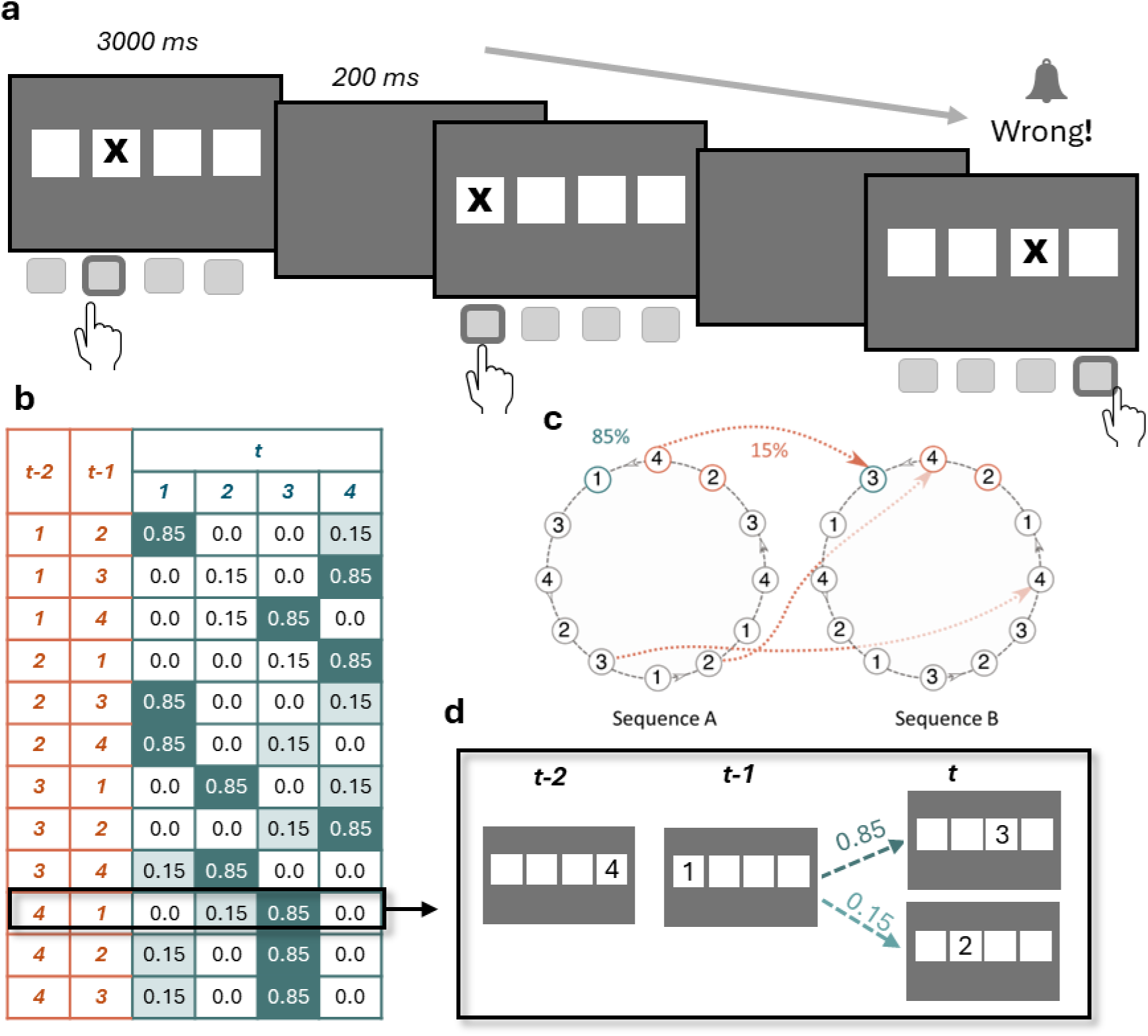
Probabilistic Serial Reaction Time task. A) Schematic representation of the stimuli and paradigm. Participants were instructed to indicate the location of the stimulus by means of a button press corresponding to each of the 4 possible locations. The task consisted of two sessions (Pre- and Post-reversal), comprising 960 trials each. B) Transition probabilities of the stimulus locations. The location of the stimulus on each trial was decided by two Markov chain sequences. The two sequences differed in the second-order conditional occurrence, with one sequence occurring 85% of the time and the other 15%, leading to High and Low Probability trials. C) The two deciding sequences, with example transitions between sequences. Here, Sequence A (85%) has high probability of occurring while Sequence B has low probability of occurring (15%). Following one session (960 trials) of the task, the probabilities of each sequence occurring were reversed, leading to Sequence A as the low probable sequence and Sequence B as the high probable one. D) An example transition leading to High and Low Probability trials.

Our findings demonstrate that participants implicitly learned the probabilistic structure of the task, evidenced by the speeding of responses on high versus low probability trials as the task progressed (*Fig 2*). Post-error slowing indicated that errors played a key role in driving learning (*Fig 2*). These model-agnostic results were further supported by computational modelling (*Fig 3*) findings, which suggest that learning was driven by surprise and beliefs about volatility (*Fig 4*). Importantly, we found strong correlations between participants’ M1 glutamate + glutamine (Glx) levels and prediction errors and volatility beliefs (*Fig 5*). This suggests that M1 Glx plays a crucial role in high-level volatility beliefs during implicit learning under uncertainty. We also found that, while individuals high in trait anxiety were faster *after* the reversal, contrary to our predictions, there was no association between trait anxiety and volatility beliefs. Our findings offer novel insights into the computations of probabilistic learning and reveal important neurochemical markers of adaptive behaviour in unpredictable environments.

**Figure 2.**
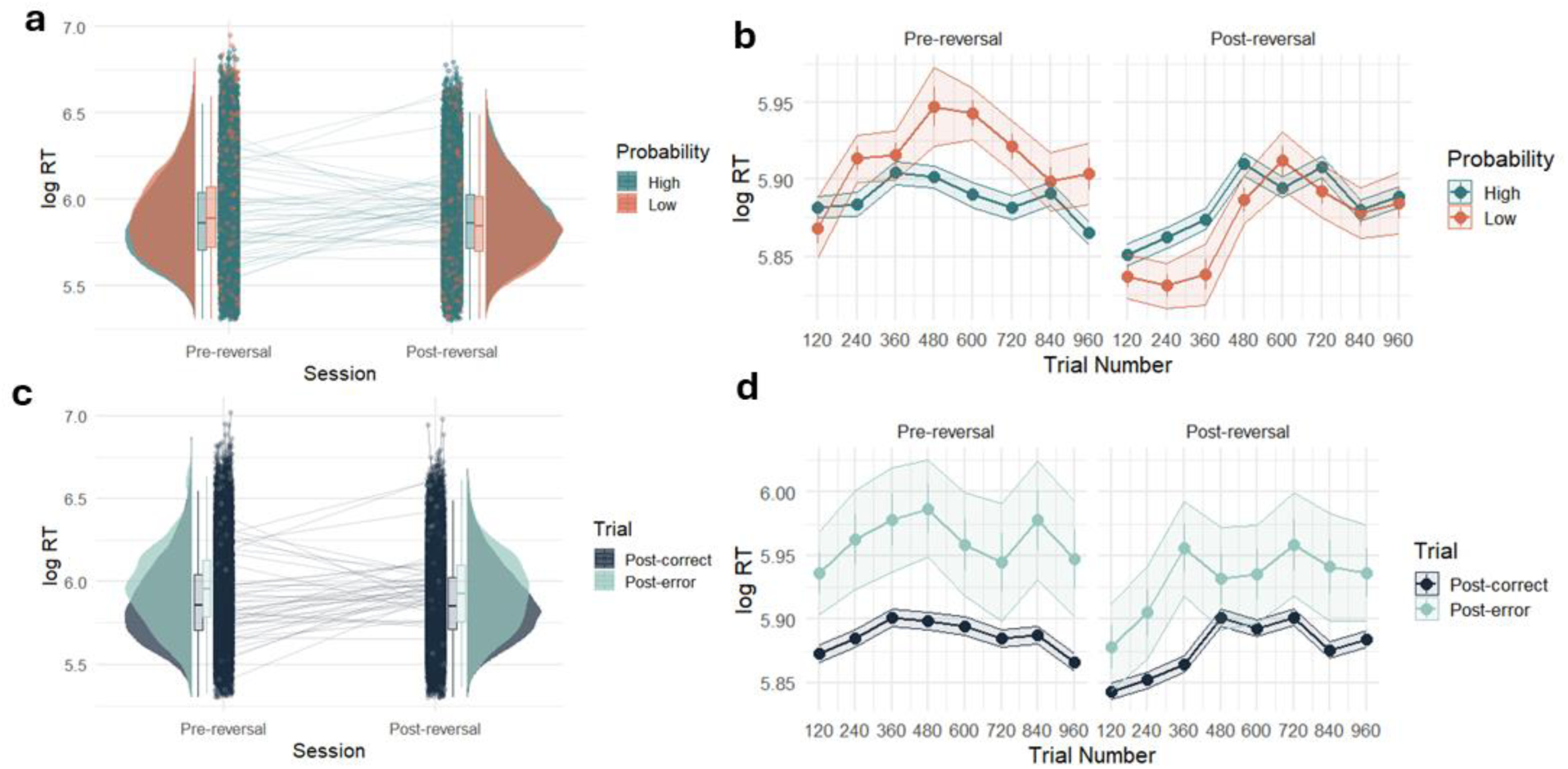
Model-agnostic behavioural results. a) Distributions (boxplots and raincloud plots) of mean log-transformed RT for correct responses grouped according to High Probability (green) and Low Probability (orange) trials across Pre- and Post-reversal sessions. Error bars of boxplots show the standard error of the mean. Dots and corresponding lines indicate individual participant means across the sessions. b) Mean log-transformed RT across time for High Probability (green) vs Low Probability (orange) trials across Pre- and Post-Reversal sessions. Data points indicate the mean for each block (120 trials), and bars indicate the SEM. c) Distributions (boxplots and raincloud plots) of mean log-transformed RT for all responses grouped according to whether trials followed an error (post-error, in turquoise) vs a correct (post-correct, in dark blue) response. Error bars of boxplots show the standard error of the mean. Dots and corresponding lines indicate individual participant means across the sessions D) Mean log-transformed RT across time for Post-error vs Post-correct trials across Pre- and Post-Reversal sessions. Data points indicate the mean for each block (120 trials), and bars indicate the SEM.

**Figure 3.**
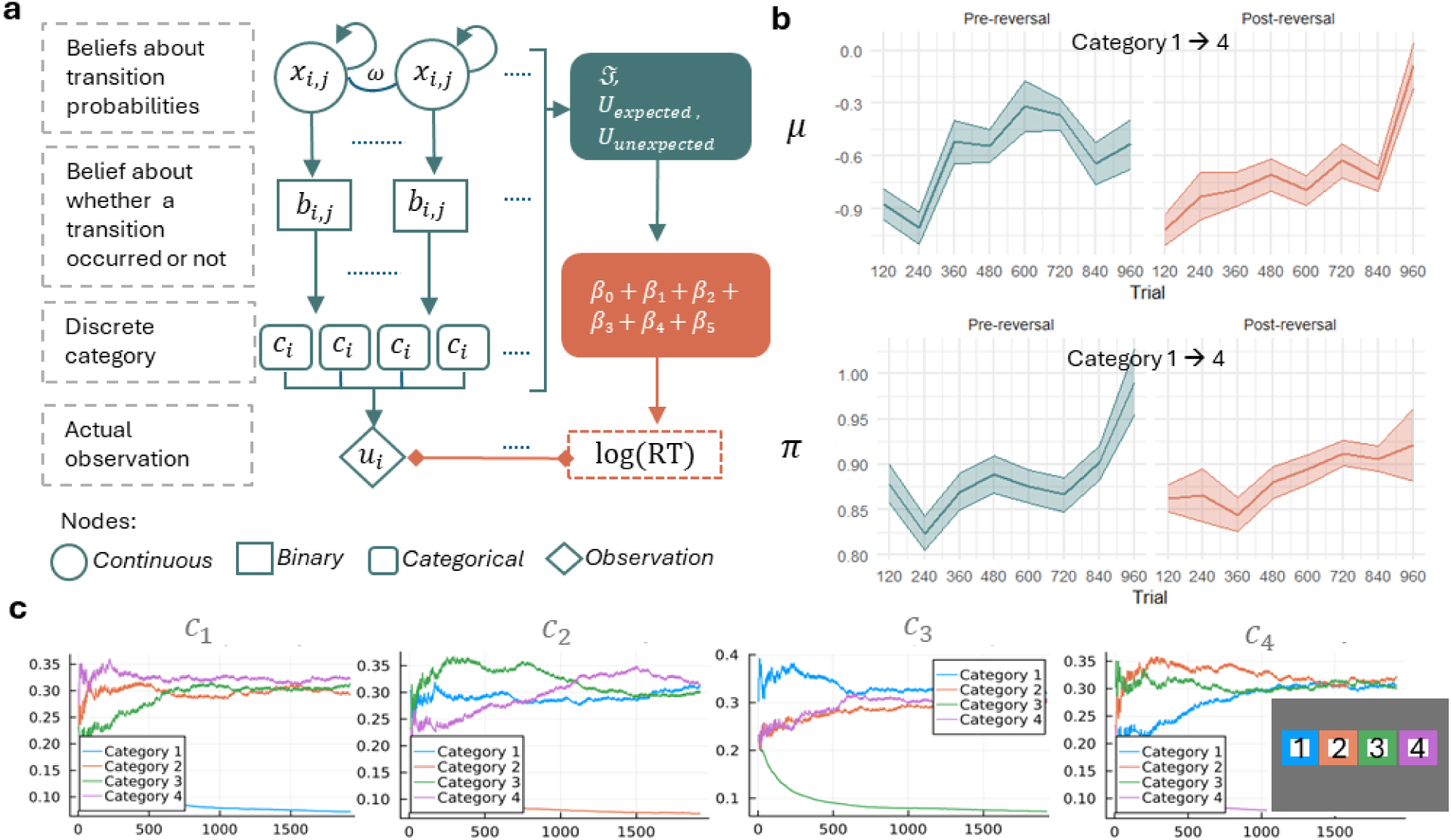
Categorical state-transition Hierarchical Gaussian Filter (HGF). a) Schematic description of the categorical state-transition HGF. Beliefs are represented at probabilistic nodes, with higher-level continuous nodes (*x*_*i*,*j*_) predicting the changing transition probabilities represented by lower-level nodes (*b*_*i*,*j*_), contingent on the belief about general environmental volatility (ω) that is shared between all x_i,j_. Transition probabilities are normalised to proper probability distributions in categorical nodes c_i_. Straight arrows indicate dependencies in the HGF’s generative model, and curved arrows indicate a dependency on the node’s own previous time step. Dotted lines indicate the multiplicity of nodes. Key belief states from the HGF are mapped to reaction times by means of a response model (indicated in orange). b) Evolution of higher-level beliefs µ (top graph) about transition probabilities and the corresponding posterior precision *π* (bottom graph) at the continuous node (*x*_*i*,*j*_) for a specific transition (namely, from Category 1 at trial t-1 to Category 4 at trial t), at an example session. Each data point corresponds to the mean of the state transition belief (µ) or posterior precision (*π*) in bins of 120 trials (x-axis), across each session of the task (Pre-reversal in green, and Post-reversal in orange). c) The HGF’s inferred state-transition probabilities over time in the same example session. Each panel represents transitions from one of the 4 discrete categories (i.e., possible stimulus locations). Transition beliefs (y-axis) across time (x-axis) are estimated for each possible transition. For example, the leftmost panel (titled c1) represents the transitions from Category 1 (blue) to: Category 1 (blue), Category 2 (orange), Category 3 (green), and Category 4 (purple). As the task does not contain transitions from Category 1 to Category 1, the estimate of this transition quickly approaches 0.

**Figure 4.**
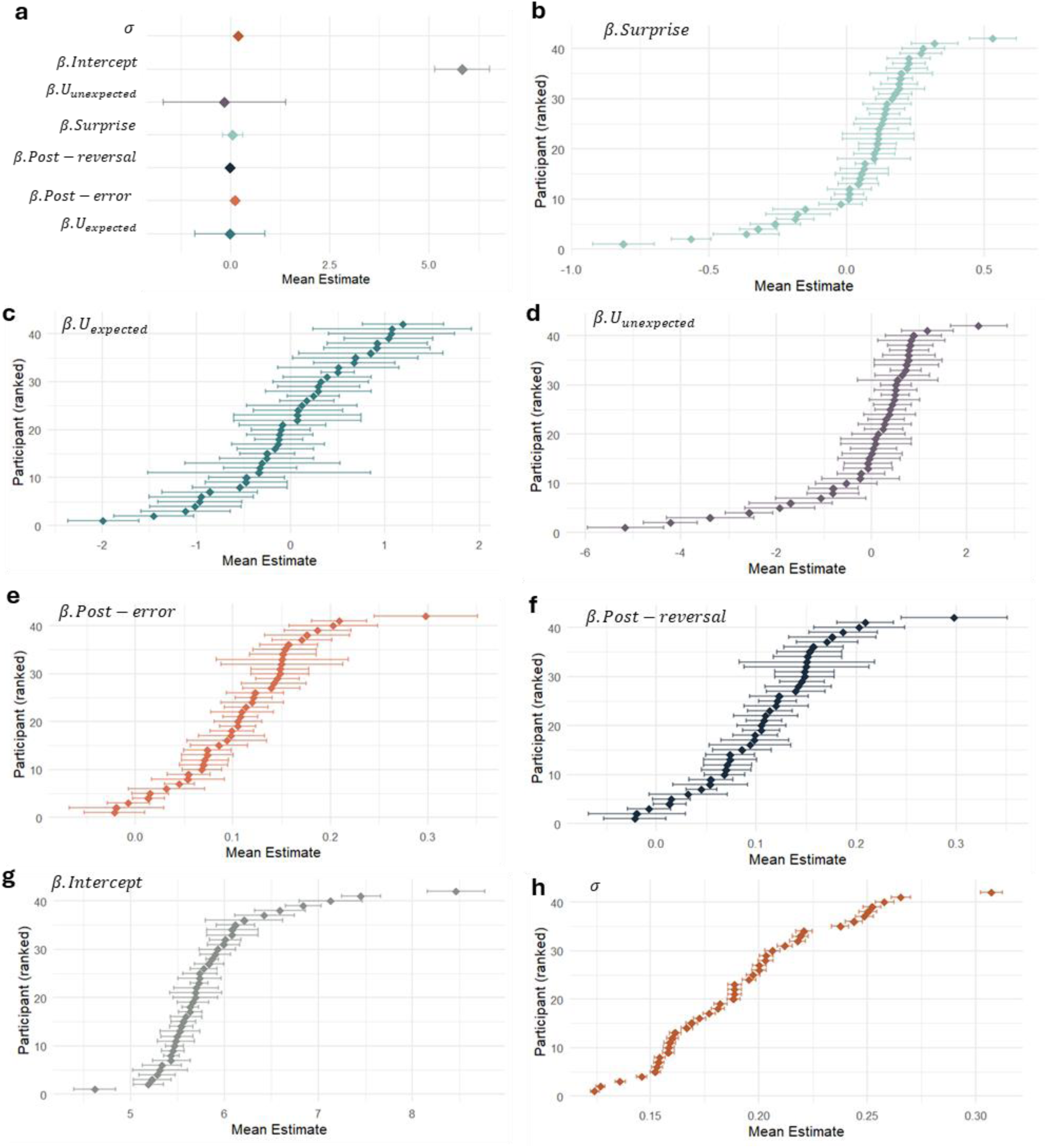
Categorical state-transition HGF results. a) Mean beta weights at the group level for each of the predictors included in the response model. The HGF response model maps beliefs estimated from the perceptual model and relevant task regressors onto participants’ log-transformed RT by means of a linear regression. Data points indicate the mean estimates (x-axis), with bars representing SEM. All the predictors (y-axis) were included in the same response model. b-h) Individual participant-level mean beta estimates for response model predictors *β*. Surprise (pale green), *β*. (dark green), *β*. *U*_*unexpected*_ (purple), *β*. *Post* − *error* (orange), *β*. *Post* − *reversal* (dark blue), *β*. *Intercept* (grey), and standard error of the regression (*σ*) (brown) respectively. The x-axis indicates the mean estimate and the y-axis indicates each participant arranged in ascending order of the value of their individual mean estimate. Each data point corresponds to the participant’s mean, with error bars representing SEM. Posteriors were estimated through Monte Carlo Markov Chain (MCMC) inference using 4 chains and 2,000 iterations corresponding to each chain.

**Figure 5.**
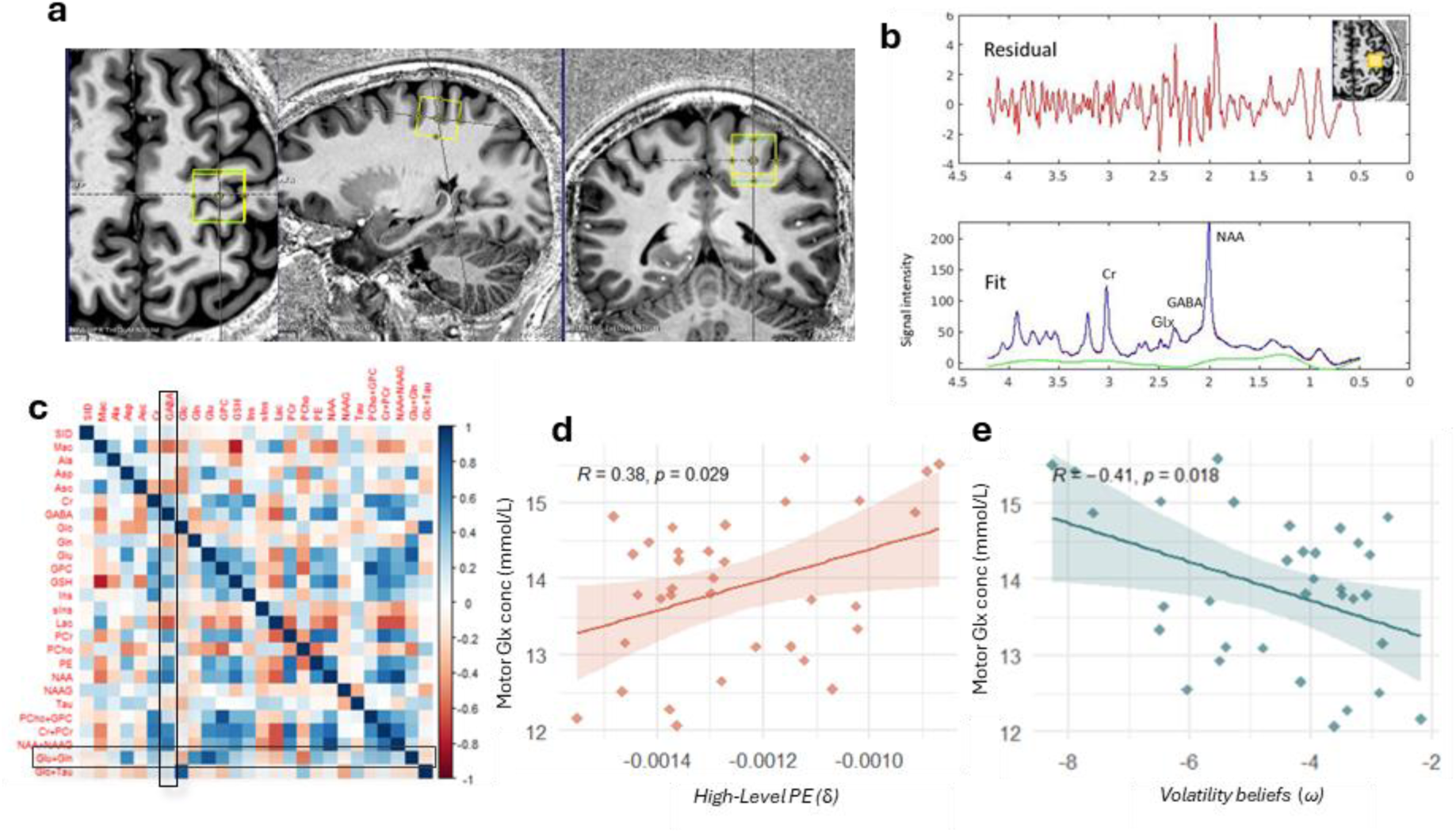
MRS spectroscopy. a) Motor cortex (M1) voxel placement. b) Example MR spectra for a sample participant. Upper panel: raw spectra containing the residuals after LC Model fitting; Lower panel: LC Model fit with peaks of all metabolites. The main Glx (Glu + Gln) peak resonates between 2.2 and 2.4 ppm, while GABA shows peaks at approximately 3.0ppm, 1.9ppm and 2.2ppm. c) Correlation matrix of all the quantified metabolites (uncorrected) in M1 voxel across all participants. The metabolites GABA and Glx (Glu + Gln) are outlined. Colour bar indicates the strength of the correlation, with dark blue indicating positive correlation and red indicating negative correlation. SID refers to Subject ID. d) Significant positive correlation between M1 Glx levels and high-level prediction errors (r = 0.38, p=0.02) for n=37 participants, e) Significant negative correlation between M1 Glx levels and predicted belief about volatility (r= -.41, p =0.01) for n=37 participants.

## Results

### Implicitly-learned probabilities and errors drive task performance

We implemented a probabilistic serial reaction time (SRT) task in which participants were told to indicate the location of a stimulus by pressing keys corresponding to four possible locations (*Fig 1a*). Unbeknownst to the participants, the stimulus location on each trial was associated with pre-determined probabilities that may be implicitly learnt over time (*Fig 1b-d*, *Methods*). Following one session of the task, there was a switch to the probabilities to gauge the impact of a probabilistic reversal on implicit learning.

To further examine whether the probabilistic sequences were implicitly learned, we fit linear-mixed effects (LME) models to the data. We examined the main effects of stimulus probability (High vs Low Probability) and session (Pre-vs Post-reversal) on log-transformed RT (*LME model 1*). This indicated main effects of both stimulus probability (*b*= 0.004, *SE*=0.001, *t*(79,200)=2.48, *p*=0.013) and session (*b* =-0.009, *SE*=0.001, *t*(79,200)=−7.4*, p* < 1.28×10^−13^) (*Fig 2a*).

To examine the effect of learning over time, we created three bins of 320 trials each, corresponding to early, middle, and late stages of learning. The binning was done separately for each session of the task. To test for interactions, earning stage was included as a predictor to the above model (*LME Model 2*). We found a significant interaction between stimulus probability and learning stage; specifically, for low probability stimuli, reaction times were significantly slower in the middle stages compared to the early stages (*b*=0.023, *SE* = 0.004, *t* = 5.13, *p* = 2.86×10^-7^), while they were significant faster for late stages compared to early stages (*b*= 0.020, *SE* = 0.004, *t* = 4.57, *p* = 4.74×10^-6^). The interaction between session and stage was also significant. In the post-reversal session, participants were faster in the *early* stage compared to the middle stage, suggesting a possible increase in learning in response to the volatility induction (*b*= 0.026, *SE* = 0.003, *t* = 8.28, *p* < 2×10^-16^). The responses sped up during the late stage of the post-reversal session (*b*=0.041, *SE* = 0.003, *t* = 12.69, *p* < 2×10^-16^). A more detailed visualization of the temporal effects can be found in *Fig 2b*.

Overall, accuracy rates were high across the pre-reversal (*Mean*= 0.92, *SD*= 0.20) and post-reversal (Mean=*0.93, SD=* 0.20) sessions. We examined whether trial-by-trial errors had an effect on probabilistic learning by implementing an LME model on log -transformed RT with outcome of the previous trial (correct vs wrong) and session (pre-vs post-reversal) as predictors (*LME Model 3*). We found a main effect of errors on response speed of subsequent trials (*b* = 0.11, *SE* = 0.008, *t*(79,200)=13.15, *p* < 2 x 10^-16^). In addition, we found an interaction between post-error trial and session (*b*= −1.67×10^2^, *SE* = 7.9×10^-3^, *t*=2.11, *p* = 0.03), suggesting a reduced effect of post-error slowing in the post-reversal session (*Fig 2 c-d*).

### Categorical state-transition HGF

Using Bayesian inference approximated by Markov Chain Monte Carlo (MCMC) methods, the complete task dataset was fit to three different learning models of which categorical state-transition HGF performed best on model comparisons (*Supplement*). The categorical state-transition HGF learns transition probabilities between observed discrete categories using variational Bayesian inference in a predictive coding-like manner (*Fig 3a*, *Methods*). Here the categories tracked by the HGF represent the four possible stimulus locations on the SRT, resulting in a total of 16 possible transitions. The HGF is implemented as a network of nodes, each representing a belief about some part of the task: the received sensory observation *u* of category *j*, the transition from the previous category *i* that was observed *c_i,j_,* the probability of that transition occurring on the next trial *b_i,j_*, and the in-time change of the probability in log-odds *x_i,j_*. On each trial, the observed category is compared to its prediction to produce a measure of surprise ℑ (*Equation 1*) and a prediction error δ *(Supplement)*. These quantities are passed up through the hierarchy of nodes and used to update beliefs about transition probabilities, which in turn supply predictions for the following trial (*Methods, Supplement*).

The HGF holds Gaussian beliefs about *x_i,j_*, with the mean *μ* representing the current belief about the transition probability and the precision *π* representing the certainty in that belief (see *Fig 3b-c* for an example of how these two values evolve over time for a specific transition probability). These beliefs are updated faster when prediction errors are large, and when the precision *π* of the current belief is low. At each trial, the means *μ* of the belief are transformed to probability space and normalised to construct the final expected transition probabilities (*Fig 3d* displays an example session where these 16 transition probabilities are learnt over time). The learning of transition probabilities is contingent on the parameter *ω*, which represents the expectation of general volatility in the environment. Higher values of *ω* lead to less certainty and faster updating of beliefs.

To connect the belief dynamics of the HGF to the log-transformed reaction times, ℑ (*Equation* 1), *U*_*expected*_ (*Equation* 2), *U*_*unexpected*_ (*Equation* 3), *Post* − *error*, and *Post* − *reversal* were included as predictors in a response model (*Equation 5)*. Thus the full model has 8 free parameters: the expectation of general environmental volatility *ω*, the 6 beta parameters (including intercept term) of the response model, and the standard error of the regression σ.

At the group level, the mean and SD estimate for each regressor of the response model were as follows: *β*_0_ (5.83,0.68), *β*_1_. ℑ(0.053, 0.25), *β*_2_. *U*_*expected*_(−0.02, 0.89), *β*_3_. *U*_*unexpected*_(− 0.15, 1.5), *β*_4_. *Post* − *error*(0.10, 0.07), and *β*_5_. *Post* − *reversal*(−0.01, 0.05) (*Fig 4a*). To test if there were consistent effects across the whole group, we conducted one-sample *t*-tests separately for each beta estimate. We found that only *β*_4_. *Post* − *error* had a significant effect (t(41)=10.739, 95% CI [0.08,0.12], *p*<0.001), indicating that the post-error slowing effect was most stably present across the group. However, at the individual-level, we see clear indication of inter-individual variation in the distributions of estimates for each regressor of the response model (*Fig 4 b-h*).

### M1 Glx predicts probabilistic reversal learning through prediction errors and beliefs about volatility

We fit an LME model on log RT with task session (Pre vs post -reversal) and metabolite levels as regressors (*LME Model 4*). We found a significant interaction between M1 Glx and session (*b* = 0.039, *SE*= 0.001, *t*(64,470)= 2.68, *p* = 0.007), suggesting that M1 Glx may affect *reversal* learning in particular. Next, we conducted separate Pearson’s correlations between the computational model estimates and metabolite levels. To calculate each participant’s mean prediction error (δ), high-level prediction errors (δ) corresponding to the 16 possible transitions were averaged for each participant and included in the correlation.

This yielded a significant positive correlation (*r*(32)=0.37, *95%CI*[0.04,0.63], *p*=0.029). In other words, higher levels of M1 Glx were associated with larger prediction errors (δ). We also found a significant negative correlation between M1 Glx levels and participants mean belief about volatility (*ω*) (*r*(31)=−0.40, *95%CI*[−0.659,−0.075, *p*=0.018), indicating that higher Glx levels are associated with lower beliefs about volatility. To confirm the regional- and metabolite-specificity of our findings, we ran supplemental control analyses on M1 GABA and a control voxel placed on the occipital cortex. We found no relationships between M1 GABA and behaviour, confirming that these findings are specific to Glx (*Supplement*). In addition, we examined Glx levels from MRS data acquired from a control voxel; this yielded no significant results (*Supplement*).

### Relationship with Trait Anxiety

We implemented an LME model on log-transformed RT with task session and trait anxiety scores as predictors, and state anxiety as a controlling variable (*LME Model 5*). This yielded a significant interaction between trait anxiety and task session (*b* = −1.9, *SE* = 0.0001, t(79) = - 15, *p* = 2×10^-16^). In other words, individuals high in trait anxiety, although overall slower compared to those with low trait anxiety, were found to be faster after the reversal. To visualise this effect, we created High and Low Trait Anxiety groups based on a median split of the trait anxiety scores (*Fig 6a*).

**Figure 6.**
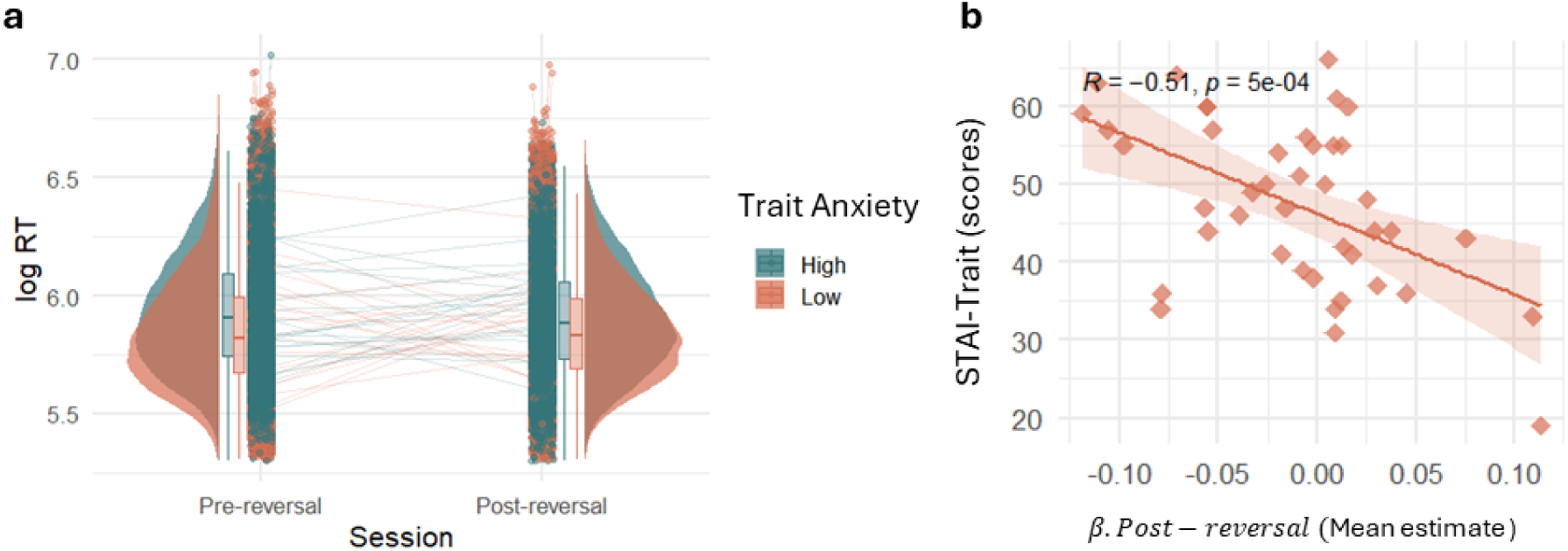
Relationship with Trait Anxiety. a) Distributions (boxplots and raincloud plots) of mean log-transformed RT grouped according to High Anxiety (green) versus Low Anxiety (orange) across Pre- and Post-reversal sessions. Error bars of boxplots show the standard error of the mean. Dots and corresponding lines indicate individual participant means across the sessions. Anxiety levels were split by median trait anxiety scores for visualisation only and analyses were focused on continuous trait anxiety scores. B) Significant negative correlation between HGF response model beta estimate for post-reversal and trait anxiety scores for 42 participants (r= −.51, p = 5 x 10^-4^).

Next, to corroborate the above model-agnostic findings, we examined relationship between trait anxiety (as calculated by STAI-Trait scores) and the HGF response model beta estimate for task session *β*. *Post* − *reversal*. This yielded a significant negative correlation (*r* = - 0.51, t(42) = −3.78, *95% CI* [-0.70, −0.24], *p* = 5 x 10^-4^), suggesting that the magnitude of the relationship between response speed and the reversal may be predicted by trait anxiety scores (*Fig 6b*). Contrary to our hypothesis, we found no relationships between beliefs about volatility (*ω*) and trait anxiety (*r* = −.17, t(40)= −1.06, 95% CI [-0.44 0.14], *p*=0.29).

## Discussion

This study provides evidence of motor glutamate + glutamine (Glx) contribution to probabilistic learning in humans, specifically in environments that involve changing or volatile statistics. The hierarchical Bayesian model we implemented offers a robust framework for understanding how individuals build and update beliefs about the world, capturing both the explicit and implicit processes involved in learning under uncertainty and the complex belief trajectories of non-binary outcomes. Our findings suggest that variations in M1 Glx may serve as a neural signature of individual differences in adaptive learning under uncertainty.

We found that participants implicitly learned the probabilistic structure of a four-choice sensorimotor task and adapted to changes in volatility induced by a probabilistic reversal. Evidence of post-error slowing suggested that participants adjusted their behaviour following errors to improve subsequent performance. Additionally, the reversal of probabilities led to faster responses in the early and late stages of the post-reversal session, indicating that individuals adapted to the new contingencies. The computational modelling findings confirmed these model-free results. At the group-level, post-error slowing most consistently contributed to reaction times during probabilistic reversal learning. Notably, at the participant-level, we see inter-individual differences in the response model beta estimates, including surprise, expected and unexpected uncertainty, and post-reversal learning, revealing distinct “uncertainty fingerprints” (*Fig 4*). We further linked the model-free and model-based results by means of a supplemental analysis where computationally-estimated surprise was matched against the trial-by-trial behavioural data. We found that the temporal evolution of surprise aligned with our model-free behavioural findings (*Supplement*). These results add to the growing body of literature involving different populations, including recent work disentangling volatility from stochasticity^51^, demonstrating that volatility enhances – rather than impedes – learning ^5,10,12,26–28,52^.

Based on prior findings linking anxiety and related conditions to misestimation of uncertainty and overestimation of volatility ^9,12,17,20,28^, we had hypothesised a relationship between trait anxiety levels and beliefs about volatility (*ω*). In contrast, we found no association between trait anxiety and volatility beliefs (*ω*) in our task, suggesting that individuals with high trait anxiety may not be as negatively affected by uncertainty in implicit probabilistic learning tasks that lack valence-related elements (such as reward, punishment, or threat). Notably, although individuals with high trait anxiety were generally slower or more cautious, they responded more quickly after the reversal, suggesting that they effectively adapted to the changed contingencies. This is line with recent findings that individuals high in trait anxiety may be faster to update their expectations following reversals^19^.

Our key contribution lies in linking inter-individual differences in processing uncertainty — "uncertainty fingerprints" — to neurochemical levels in humans. We found a significant interaction between M1 Glx and task session, suggesting that Glx in the primary motor cortex plays a role in reversal learning on a probabilistic sensorimotor task. Numerous studies on non-human primates and rodents have pinpointed the neural basis of reversal learning to glutamatergic mechanisms ^44,45,53–58^. We establish a similar link in humans and delve deeper into this by means of computational modelling. Our model-based findings suggest that higher M1 Glx levels are associated with greater prediction errors, reinforcing the idea that glutamate-driven plasticity supports error-driven learning. Notably, we also observed a negative correlation between Glx levels and beliefs about volatility, indicating that participants with higher M1 Glx levels might have lower expectations of general volatility in the environment. These results are consistent with the idea that Glx may be crucial for processing uncertainty and updating beliefs about the environment. In contrast, we found no significant relationships between M1 GABA and uncertainty processing, confirming the glutamatergic specificity of these findings. The absence of a relationship between GABA and probabilistic learning in our study suggests that the excitatory actions of Glx, rather than inhibitory regulation via GABA, are more central to the dynamics of learning in this task.

While the present study provides valuable insights into the role of Glx in learning under uncertainty, there are a number limitations that should be considered. First, baseline MRS measurements may be considered *indirectly* indicative of neurotransmitter signalling. Furthermore, as our MRS analyses were focussed on the Glx complex, which contains signals from both glutamate and glutamine, we cannot attribute our findings to glutamate in particular. Glutamine is a precursor to glutamate and the two are enzymatically linked ^59^. We were able to confirm the regional specificity of our findings based on the results of a control analysis conducted on an occipital cortex voxel. However, other brain regions and neurotransmitter systems likely play significant roles in learning and decision-making under uncertainty. Future studies incorporating multi-region MRS imaging or other neuroimaging techniques (such as fMRI) could provide a more comprehensive understanding of the neurobiological mechanisms underlying uncertainty processing in humans. Additionally, examining excitatory mechanisms in clinical populations with anxiety or other psychiatric conditions could provide important insights into how neurochemical imbalances contribute to maladaptive learning and decision-making.

In conclusion, this study presents a novel investigation into the neural basis of probabilistic learning under uncertainty, revealing that M1 Glx plays a critical role in beliefs about volatility and prediction errors during implicit learning. We further present a novel model of probabilistic learning designed to capture the complexity of non-binary choices in the classic four choice probabilistic Serial Reaction Time task. By linking computational estimates of uncertainty processing with neurochemical data, we provide a deeper understanding inter-individual differences in adaptive behaviour in unpredictable environments. These results also offer potential avenues for future research into how neurochemical variations in M1 Glx may influence learning and decision-making in both healthy and clinical populations.

## Methods

### Participants

43 right-handed participants (20 Female: 23 Male), aged 19-39 years (*Mean*=28.37, *SD*= 4.75) participated in the study. Right-handed participants with normal or corrected-to-normal vision were recruited through social media, local classified advertisements, and University mailing lists. The sample size was based on a power calculation using effect sizes from recent literature ^60–63^. To estimate a correlation of *r*=0.4, a sample size of n=42 (power = 0.8, Type I error = 0.05) was required to observe our primary effect of interest, namely a relationship between the metabolite of interest and behaviour.

### Procedure

This study was approved by and conducted in accordance with the regulations of the University of Cambridge Psychology Research Ethics Committee (PRE.2020.127). Written informed consent was obtained from all participants. The study was completed over two sessions.

In the first session, pparticipants completed the probabilistic SRT task ^3,64^ on a desktop computer at a viewing distance of 50 cm from the screen in a darkened room. Stimuli were presented using *Psychtoolbox version 3* ^65^ in *MATLAB R2019b*. Stimuli were displayed on a 24” monitor at a resolution of 1920×1080. Each trial consisted of the appearance of a cross in one of the 4 squares (*Fig 4a*). Participants were instructed to indicate the location of the cross by pressing a key corresponding to the four different possible locations. Stimulus duration lasted 3000 *ms* with an inter-stimulus interval of 200 *ms*. A beep was played to alert participants to incorrect or missed responses. Following the first session, participants commenced the second session after a 15 minute break. The probabilistic version of the SRT task differs from the deterministic version in its use of second-order conditional sequences ^66,67^. Here, unbeknownst to the participants, the position of the stimulus was determined by one of two Markov-chain sequences (*Fig 1c*):

Sequence A: 1-2-1-4-3-2-4-1-3-4-2-3-1-2-1-4-3-2-4-1-3-4-2-3
Sequence B: 1-3-2-3-4-1-2-4-3-1-4-2-1-3-2-3-4-1-2-4-3-1-4-2

The position on trial *t* was determined by the position of the two previous trials, *t-1* and *t-2*. The two deciding sequences differed in the second-order conditional occurrence, with one occurring 85% of the time and the other 15% ^67–69^ (*Fig 1b-d*). In the first session, there is an 85% probability of trial *t-1* and *t-2* leading to trial *t* in Sequence A, and a 15% probability of transitioning to trial *t* in Sequence B. Participants are hypothesised to develop expectations of upcoming targets in the probable sequences by implicitly learning this second-order conditional information. Following the completion of the first session, the sequences were reversed, making Sequence B is the more probable sequence, and Sequence B the less probable sequence.

In the same session, participants also completed the Spielberger State-Trait Inventory (STAI), a 40-item questionnaire to assess individuals’ anxiety across state and trait domains (Spielberger, 2010).

In the next session, participants completed a 7T MRI scan at the Wolfson Brain Imaging Centre, University of Cambridge. A Siemens 7T Magnetom Terra scanner (Siemens, Erlangen, Germany) with a single-channel transmit and 32-channel receiver head coil (Nova Medical, Carson, CA) was used. T1-weighted MP2RAGE structural scans (repetition time = 4300 ms, echo time = 1.99 ms,, bandwidth= 250 Hz/px, voxel size = 0.75 mm^3^, isotropic field of view = 240 x240×157 mm, acceleration factor = 3, flipangle = 5/6° and inversion times = 840/2370 ms) were acquired. Spectra were acquired using a short-echo semi-LASER sequence ^70,71^ with repetition time/echo time = 5000/26 ms, 64 repetitions. Pre-scan optimisation included FASTESTMAP B_0_-shimming ^72^, unsuppressed water-peak seriesB_1_ calibration, and VAPOR water suppression calibration ^73^. MR spectra were obtained from a 2×2×2 cm^3^ voxel of interest (VOI) placed on the primary motor cortex (M1) while participants were at rest. The voxel placement was manually centred on the omega- or epsilon-shaped left hand knob area using the central sulcus as a landmark ^61,74^ (*Fig 5a*). The VOI was placed to capture the entire hand knob and to exclude the dura. For the motor VOI, the anatomical landmarks could be most clearly observed from the coronal view (*Fig 5a*). To confirm the regional specificity of the findings, we also obtained MR spectra from a control voxel placed in the occipital cortex, acquired with identical parameters (*Supplement*).

### Analyses

Data were cleaned and analysed using *R 4.3.3*. Behavioural data outliers were identified as RT < *200 ms,* while slow RT outliers were computed separately for each participant as trials with RT more than 2 standard deviations from their overall RT. We employed linear mixed-effects models (LME) to examine the effect of task, questionnaire, or neural variables on log-transformed RT using R packages *lme4* ^75^ and *lmerTest* ^76^. This approach allows us to account for both fixed effects (variables that were consistent across all participants) and random effects (variables that vary across participant). Each participant was included as a random intercept in the LME models. LME models testing for effects of stimulus probability (*LME Models 1 and 2*, *Results, Fig 2*) on response time were focused on correct responses only. The following LME models were employed:

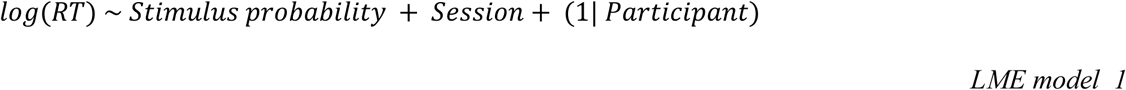

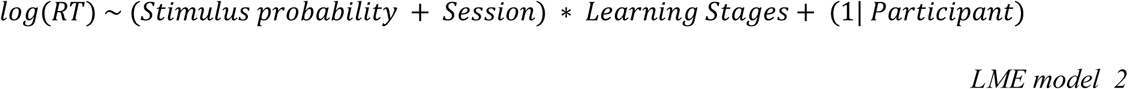

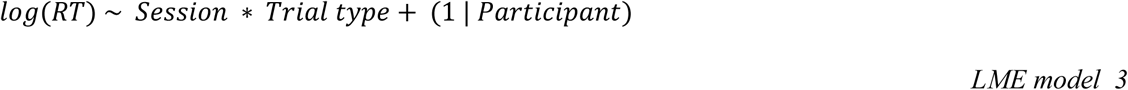

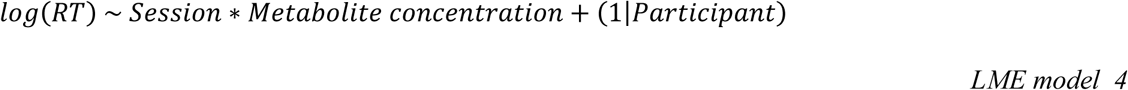

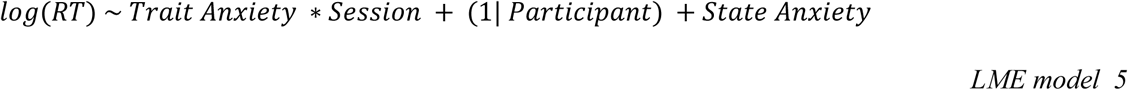

where stimulus probability refers to high vs low probability trials, session refers to pre-vs post-reversal task sessions, trial type refers to post-error vs post-correct trials, Trait Anxiety indicates STAI-Trait subdomain scores, and State Anxiety indicates STAI-State subdomain scores.

The acquired MR spectra were corrected for eddy current effects, and phase and frequency drifts using MRspa (www.cmrr.umn.edu/downloads/mrspa/), in line with the current best-practice recommendations for pre-processing ^77^. Metabolites between 0.5 and 4.2 ppm – including Glx and GABA-were quantified using *LCModel v6.31* ^78^. Metabolite concentrations are reported with reference to the water signal (sometimes referred to as the “absolute” concentration) ^79^. They were corrected for inter-individual differences in tissue volume. Details of the tissue segmentation and correction procedure can be found in the Supplement. The following exclusion criteria were used for quality control: water linewidth > 15 Hz, signal-to-noise ratio < 40, and Cramér-Rao lower bounds > 2 standard deviations from the group median ^80–82^. Following the quality control exclusions, data from 37 participants remained for the M1 voxel. Outliers were identified as those above or below the cutoff of 2 standard deviations from the group median.

Specific relationships between the computational modelling estimates and metabolite concentrations or questionnaire measures were assessed means of Pearson’s correlations. All statistical analyses were two-tailed unless otherwise stated.

### Computational cognitive modelling

We modelled behaviour using a categorical state-transition Hierarchical Gaussian Filter (HGF). The HGF is a popular model of perception and learning in volatile environments (Mathys et al., 2011; Mathys et al., 2014) that has recently been generalized to a flexible network formalization ^36^. It is a variational Bayesian predictive processing model, where perception is modelled as a process of variational Bayesian inference on the otherwise not directly accessible states of the environment. This type of inference employs a generative model of how the environment changes, and how it generates sensory observations - the model is inverted with Bayesian inference to estimate the current state of the environment, given some sensory observation. In the generalized HGF, the generative model is implemented as a network of nodes representing different states of the environment. States can either be binary, categorical or continuous and can connect to each with *value couplings*, where the expected *value* of a node depends on another, or with *volatility couplings*, where the *noisiness* of each node depends on another. Each node type has its own set of update equations and free parameters, dependent on how they are connected. A detailed formal introduction can be found Weber et al. (2024). Here we present the updates relevant specific to our study.

We built upon the generalized HGF to create a categorical state-transition HGF, which specifically learns the probabilities of different transitions between categorical states. It consists of the following nodes: a categorical input node *u* which registers the categorical input in the task (i.e., the stimulus location); a set of categorical state nodes *c_i_* that store the type of transition made, and the normalized predicted transition probabilities; a set of binary nodes *b_i,j_* for each transition type that stores whether that transition occurred, and the predicted probability of it occurring in the future; and a set of continuous nodes *x_i,j_* that track the changing (log)probabilities of each of those state-transitions. First, node beliefs are updated based on prediction errors. Since there is no observational uncertainty, *c_i_* is simply updated by setting the belief about the observed category to be equal to the actual category. Similarly, nodes *b_i,j_* are set to 1 when the transition occurred, and 0 when it did not. Nodes *x_i,j_* are updated as continuous value parents to binary nodes; these update equations have been reported in the Supplement.

For the response model of the HGF, we follow the approach used by Marshall et al.(2016), and implement a linear regression linking various belief states of the HGF to the log-reaction times of participant’s responses.

First, the surprise (ℑ) or unexpectedness of the observed transition at each trial *t* based on the agent’s internal model is calculated as the negative log probability of the transition, under the model’s prediction (See supplement for prediction calculation):

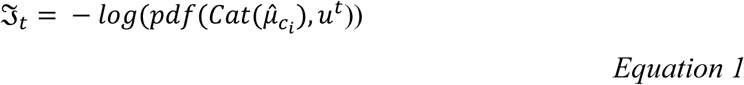

where 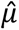 is the mean prediction, *c_i_* represents the categorical state node that stores the type of transition made, 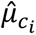 the categorical probability distribution, and *u*^*t*^ represents the node containing the actual observations at time *t*.

In addition, we quantified “expected” and “unexpected” uncertainty (Pulcu & Browning, 2019). The "expected" uncertainty is the uncertainty regarding whether the observed transition would have occurred, multiplied with the precision of the belief at *x_i,j_ :*

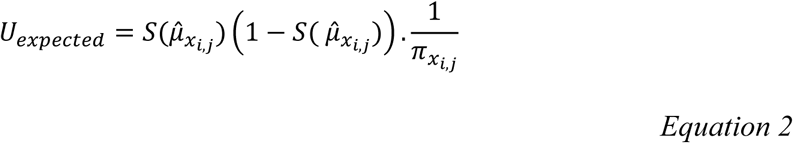

Where 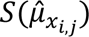 is the logistic transformation of the mean prediction for the node tracking the probability of the observed transition.

To calculate "unexpected" uncertainty, this is multiplied with the predicted volatility Ω of the transition probability, governed by the free parameter *ω*:

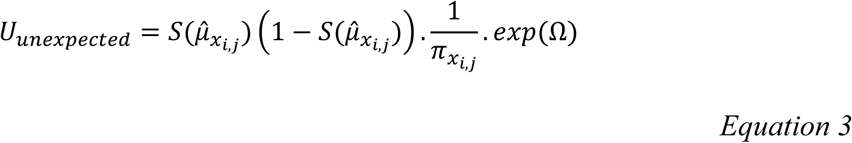

In addition to the surprise (ℑ) and the two types of uncertainty, we also included two task-related binary predictors, namely Post-error (whether the trial followed an error-trial), and Post-reversal (whether the reversal had occurred). The linear regression (including an intercept and standard error of the regression σ) equation was as follows:

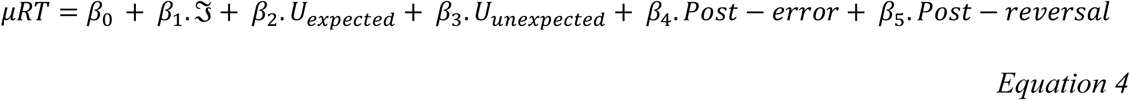

Finally, as with normal linear regression, the actual reaction times were sampled from a Gaussian *N* with standard deviation *σ* :

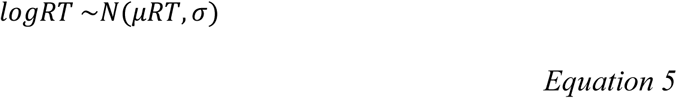

The categorical state-transition HGF with the linear regression based response model for reaction times was built and fit using the novel Julia libraries *HierarchicalGaussianFiltering.jl* v0.5.4 (https://ilabcode.github.io/HierarchicalGaussianFiltering.jl/dev/) and *ActionModels.jl v0.5.*4 (https://github.com/ilabcode/ActionModels.jl), part of the TAPAS ecosystem for computational psychiatry ^84^. Models were fit using Markov Chain Monte Carlo (MCMC) inference, specifically the NUTS() sampler ^85^ as implemented in Julia’s probabilistic modelling ecosystem *Turing.jl* ^86^. A model was fitted to the behaviour of each participant separately, using four chains with 2,000 samples each. Model convergence was confirmed through manual inspection (*Supplement, Fig S1*) and Gelman-Rubin *r̂* statistic <1.1 ^87^.

Priors were chosen based on previous research that fit the HGF to a similar probabilistic four-choice SRT task ^83^ and have been reported in the supplement (*Supplement, Table S2*). Parameter recovery was tested on a simulated dataset and has been reported in the Supplement. To assess which model best fit the data, we fit the complete dataset to three different learning models. The model comparison procedure yielded the categorical state-transition HGF as the winning model (see *Supplement* for details of model selection and comparison).

## Acknowledgements

The neuroimaging analyses and the computational models were run using resources provided by the Cambridge Service for Data Driven Discovery (CSD3) operated by the University of Cambridge Research Computing Service (www.csd3.cam.ac.uk), provided by Dell EMC and Intel using Tier-2 funding from the Engineering and Physical Sciences Research Council (capital grant EP/T022159/1), and DiRAC funding from the Science and Technology Facilities Council (www.dirac.ac.uk).

## Author contributions

NJ designed the study, acquired the funding with SBC and JS, collected the data, analysed the behavioural and neuroimaging data, visualised the task and results, ran the computational models, conducted the model comparisons, and drafted the manuscript. NJ ran the HGF models with custom Julia scripts created by PTW. PTW and CM designed the HGF model reported in the manuscript. PTW created the HGF modelling scripts, conducted the parameter recovery simulations, and contributed to the modelling sections of the manuscript. OP programmed and piloted the task and contributed ideas for behavioural analyses. FP contributed ideas for computational models, behavioural analyses, and visualisation of behavioural results, and contributed to the interpretation of results. CR and CTR designed and tested the MRI protocol. SBC and JS acquired the funding and contributed to the conception of the work. CM developed the HGF model and contributed to interpretation of results. RL supervised the overall project. All authors reviewed the manuscript.

