## Supplement for "Neurochemical markers of uncertainty processing in humans"

### MRS tissue correction

The structural T1-weighted images were pre-processed using *MATLAB R2019b* and SPM12 (www.fil.ion.ucl.ac.uk/spm/software/spm12/) using default SPM12 settings for voxel-based morphometry (VBM) and tissue segmentation [1,2]. The MP2RAGE images were first aligned to an average image in MNI space, cropped to a standard bounding box, and then segmented into tissue probability maps corresponding to grey matter (GM), white matter (WM), cerebrospinal fluid (CSF), soft tissue, bone, and air. A study-specific template was created using the DARTEL function. Images were skull stripped and warped to this template. GM and WM templates were affine transformed, warped to MNI space, and applied to each participant's tissue probability images. Tissue segmentation for each VOI was completed using the stand-alone segmentation scripts from Gannet 3.1.5 [3]. Individual MRS voxel masks were first transformed from native to MNI space and then overlaid in standard space. Metabolite levels are reported with reference to the water signal (sometimes referred to as the "absolute" concentration) [4]. We corrected the absolute metabolite concentration values for inter-individual differences in GM, WM, and CSF volumes using the alpha-correction method [5].

***Supplementary Table S1. MRS quality control metrics of final included dataset.***

| VOXEL | N | SNR Mean(SD) | LINE WIDTH *Hz* Mean(SD) | ABSOLUTE CRLB Mean(SD) | METAB-OLITE | CONCENTRATION *mmol /l* Mean(SD) | |
|---|---|---|---|---|---|---|---|
| | | | | | | Tissue-Uncorrected | Tissue-Corrected |
| **M1 (VOI)** | 37 | 79.5(15.9) | 11.7(1.1) | 0.30(0.07) | Glx | 13.4(1.08) | 14.0(1.04) |
| | | | | | GABA | 2.7(0.8) | 2.8(0.9) |
| **Occipital (Control)** | 31 | 72.5(11.4) | 13.52(0.9) | 0.30(0.05) | Glx | 13.7(1.3) | 14.1(1.2) |

**Categorical state-transition HGF update equations**

For a detailed introduction, please refer [6]). Continuous nodes $x_{i,j}$ are updated as continuous value parents to binary nodes $b_{i,j}$ via the following update equations:

$$\pi = \hat{\pi} + \frac{\kappa_{bin}^2}{\hat{\pi}_{bin}}$$

*Equation S1*

$$\mu = \hat{\mu} + \frac{\kappa_{bin}}{\pi} \delta_{bin}$$

*Equation S2*

where $\mu$ and $\pi$ are the mean and precision of the belief, respectively. $\kappa$ is the connection strength between the node and its binary child, here fixed to the default 1. $\hat{\mu}$ and $\hat{\pi}$ are the mean and the precision of the prediction respectively.

The prediction error ($\delta$) of the binary child node iscalculated as:

$$\delta_{bin} = \mu_{bin} - \hat{\mu}_{bin}$$

*Equation S3*

The predictions for the next trial are calculated, starting at the top of the hierarchy. Continuous nodes $x_{i,j}$ are calculated with the following standard equations:

$$\hat{\mu} = \lambda \mu + \rho$$

*Equation S4*

$$\hat{\pi} = \frac{1}{1/\pi_{child} + \Omega}$$

*Equation S5*

where $\lambda$ is the autoconnection parameter controlling the degree of autoregression, here set to the default of 1 (no autoregression). $\rho$ is the total predicted drift; in this model it is fixed to 0 as there is no constant drift being learned. $\pi_{child}$ is the precision of the child; as there is no observational noise, here it is infinite (i.e., the denominator only consists of the predicted volatility $\Omega$) . As there are no additional volatility parents, $\Omega$ only consists of the $\omega$ parameter - the general expected volatility of the environment. Note that, as $\omega$ is shared between all continuous nodes $x_{i,j}$,, the volatility is expected to be the same across transition types.

The prediction for the binary node only depends on the prediction of the parent (i.e., continuous nodes $x_{i,j}$ ):

$$\hat{\mu}_{bin} = \frac{1}{1 + exp(-\kappa_{bin} \hat{\mu}_{parent})}$$

*Equation S6*

$$\hat{\pi}_{bin} = \frac{1}{\hat{\mu}_{bin} \cdot (1 - \hat{\mu}_{bin})}$$

*Equation S7*

Finally, the prediction of the categorical node $c_i$ is a probability distribution consisting of the normalized predictions of all the parents tracking transitions from category $i$. Note that this normalization is necessary because of small numerical divergences due to the approximate nature of variational inference. The mean prediction of the categorical node $c_i$ is calculated as:

$$\hat{\mu}_{c_i} = \frac{\hat{\mu}_{bin\ i,j}}{\sum_j (\hat{\mu}_{bin\ i,j})} \cdots \frac{\hat{\mu}_{bin\ i,j}}{\sum_j (\hat{\mu}_{bin\ i,j})}$$

*Equation S8*

where $\hat{\mu}_{c_i}$ represents the categorical probability distribution, calculated using predictions from binary node $\hat{\mu}_{bin\ i,j}$, and $\sum_j (\hat{\mu}_{bin\ i,j})$ sums the unnormalized predictions $\hat{\mu}_{bin\ i,j}$ across all categorical transitions, resulting in a valid probability distribution.

### Model comparison

We fit the complete dataset (n=43 participants, n=1920 trials per participant) to three models that assess trial-by-trial learning. All models were fit with four chains and 2000 samples each. The categorical state-transition HGF (Model 1) is a hierarchical Bayesian model with a Gaussian process to model learning at different levels of abstraction. The Rescorla-Wagner (RW) model (Model 2) is a simple reinforcement learning model that assesses learning through prediction errors [7]. The EWA model (Model 3) is a hierarchical Bayesian learning model used to study how people adjust learning strategies based on experience; it has specifically been used in studies of probabilistic reversal learning [8–10]. The point-wise log-likelihood matrices from each model were extracted, and used to calculate Pareto Smoothed Importance Sampling (PSIS) approximation of the leave-one-out cross validation (LOO) metric for model comparison [11]. The PSIS-LOO was calculated using the "Stan" [12] and "loo" [13] libraries for R . The LOO criterion (LOOIC) values and differences in expected log predictive density (ELPD) are reported in *Table S2* . Lower LOOIC values indicates better model fit and predictive accuracy, while lower ELPD difference values indicate worse performance. Model 1 was the best-performing model, while Model 2 and Model 3 performed worse and indicated similar predictive performance given the close ELPD differences values and standard errors.

***Supplementary Table S2: Model comparison metrics***

| MODEL | LOOIC | | MODEL COMPARISON | |
|---|---|---|---|---|
| | Estimate | SE | ELPD difference | SE difference |

| | | | | | |
|---|---|---|---|---|---|
| **Model 1** | HGF | -37933.1 | 5006.9 | 0.0 | 0.0 |
| **Model 2** | RW | 97494.7 | 393.5 | -67713.9 | 2532.9 |
| **Model 3** | EWA | 97457.4 | 403.6 | -67695.2 | 2534.3 |

***Supplementary Table S3: HGF Priors***

| HGF PARAMETER | PRIORS (MEAN, SD) |
|---|---|
| $\boldsymbol{\beta_0}\ (\boldsymbol{Intercept})$ | Normal (log(500), 1.7) |
| $\boldsymbol{\beta_1}(\mathfrak{I})$ | Normal( 0, 2) |
| $\boldsymbol{\beta_2}(\boldsymbol{U_{expected}})$ | Normal (0, 2) |
| $\boldsymbol{\beta_3}(\boldsymbol{U_{unexpected}})$ | Normal (0, 2) |
| $\boldsymbol{\beta_4}(\boldsymbol{Post-error})$ | Normal (0, 1.5) |
| $\boldsymbol{\beta_5}(\boldsymbol{Post-reversal})$ | Normal (0, 1.5) |
| σ | Truncated (Normal( 0.05, 0.5), lower = 0) |
| **ω** | Normal (-3, 1) |

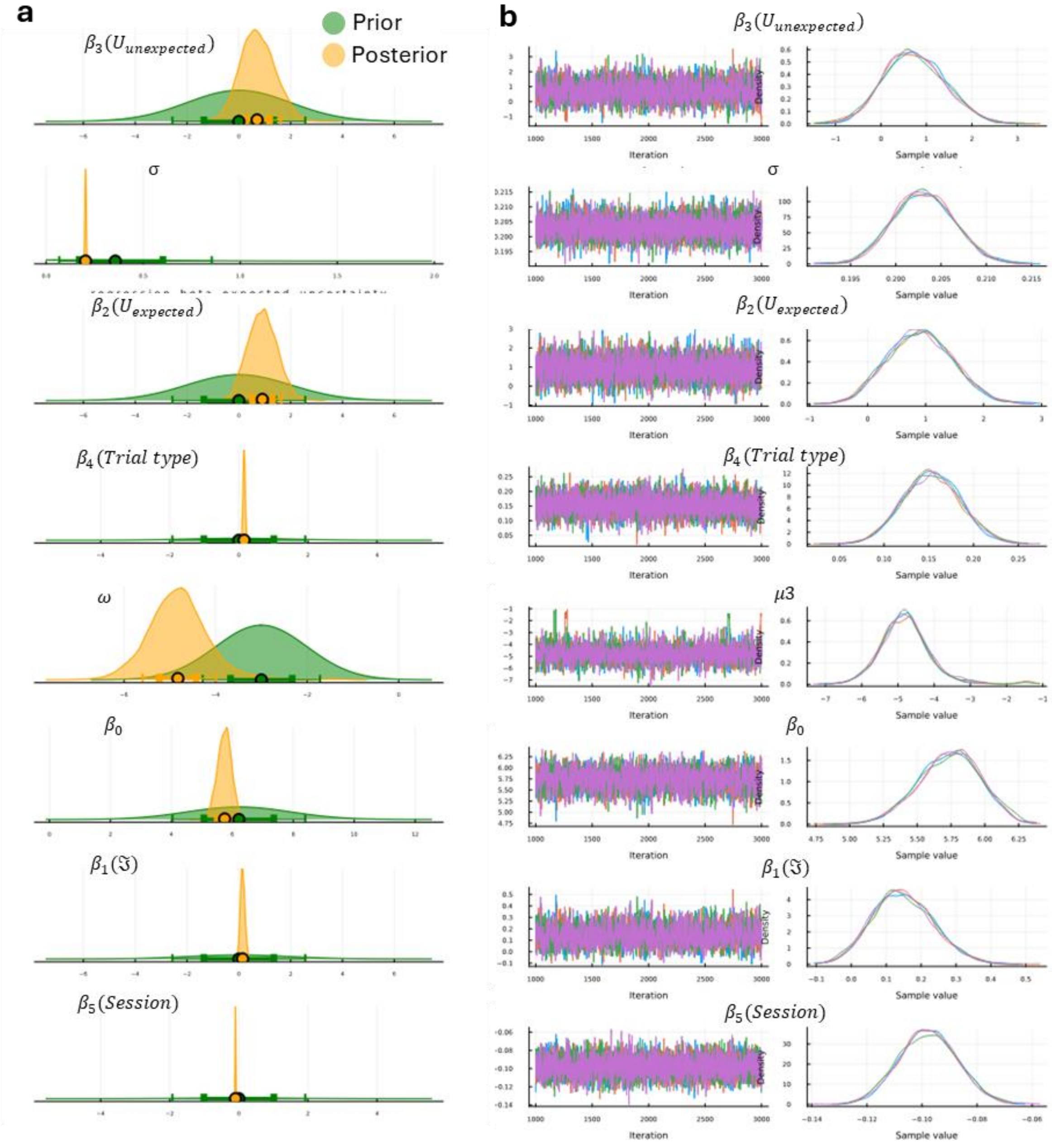


***Supplementary Figure S1. HGF model-fitting results from an example participant.*** *A) Prior and posterior probability distributions for each parameter, B) Sampling chains to indicate model convergence.*

### Evolution of surprise over time

To examine how computationally-modelled surprise ($\Im$) evolved as a function of time, we link the model-free and model-based data through an LME model with $\Im$ as the dependant variable, and session (pre- versus post-reversal), and learning stage (early, middle, late) as predictors:

$$\Im \sim Session \; * \; Learning\ stages \; + \; (1|\ Participant)$$

We found interactions between session and learning stage, with surprise increasing in the middle (b= 0.04, SE = 0.002, t = 18.90, $p < 2{\times}10^{-16}$), and late stages (b= 0.056, SE = 0.002, t

$= 25.66$, $p < 2\times10^{-16}$) of the post-reversal session. This suggests that while participants adapted to the reversal in general, their surprise fluctuated over time, in line with the model-agonistic behavioural findings.

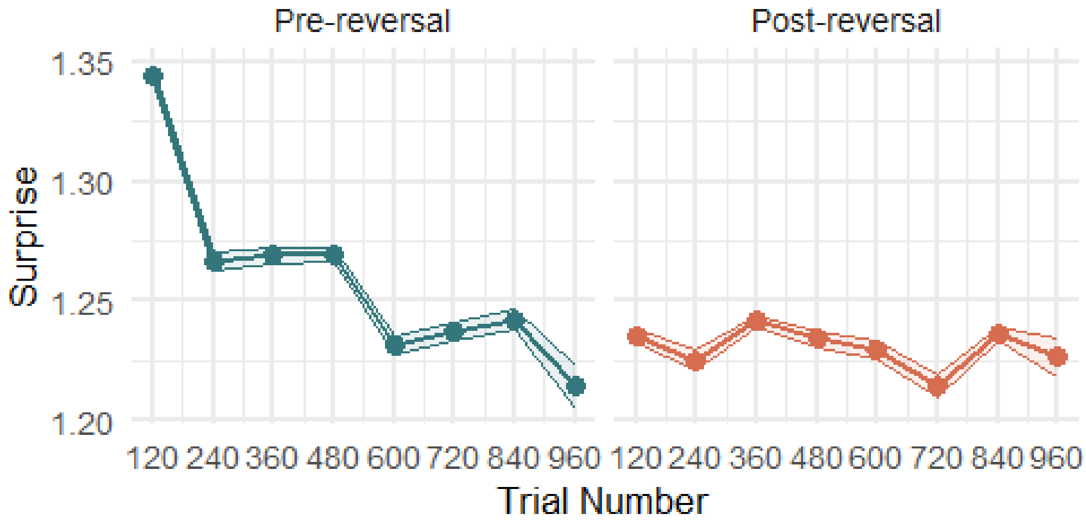


***Supplementary Figure S2. Evolution of trial-by-trial surprise ($\mathfrak{I}$ ), reflecting the unexpectedness of the categorical transition relative to the participant's beliefs****. For visualization, mean $\mathfrak{I}$ (y-axis) has been calculated for 120 trial bins (x-axis), with dots displaying the means and outlines referring to the SD. Dotted lines correspond to learning stages (early, middle, late).*

### Control MRS analyses

To test for the Glx- specificity of our findings, we ran specific linear regressions with our key variables (beliefs about volatility $\omega$ and prediction errors δ) as the dependant variable and M1 metabolites (GABA and Glx) and their respective concentrations as predictors. This yielded significant interactions between M1 Glx and a) ω ($b$ =-0.98, $SE$=0.41, $t(62)$=-2.16, $p$ = 0.03), and b) δ ($b$ = 8.6, $SE$= 4.68, t(64)=1.84, $p$=0.04). We found no evidence of any effects of M1 GABA on the same (*Fig S3 a-b*). Next, to test for the regional specificity of our findings, we repeated our main correlational analyses by replacing M1 Glx with Glx concentration acquired from the a control voxel. Here we found no correlation between control Glx levels and ω (r = -.11, 95% CI[-0.45 0.26], t(27) = -0.58, p = 0.56), and between the control Glx levels and δ (r=.14, 95% CI[-0.23 0.48], t(27) = 0.76, p = 0.45) (*Fig S1 c-d*). This confirms that Glx acquired from M1, in particular, is crucial for probabilistic reversal learning.

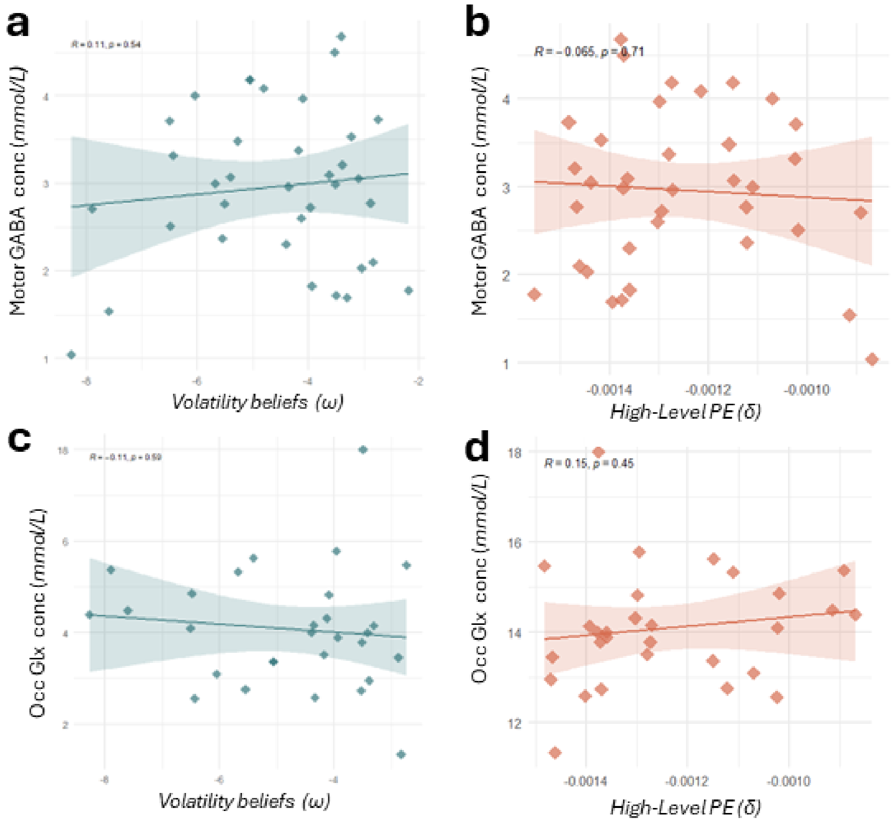


***Supplementary Figure S3. Control analyses to confirm Glx and M1 specificity of findings.*** *a) No relationship between M1 GABA and volatility beliefs (ω) (r= .11, p=0.54), b) No relationship between M1 GABA and high-level prediction errors (δ) (r = .06, p=0.71), c)No relationship between control voxel Glx and volatility beliefs(ω) (r= -.11, p =0.59 ) No relationships between control voxel Glx concentrations and high-level prediction errors (δ) (r =.15, p=0.45).*

### Parameter recovery

To test parameter recovery, we simulated synthetic behaviour with known generative parameters and fit the model to the parameter to see whether the generative parameters would be inferred. For each parameter of interest, for each of the input sequences of each of the 42 participants, and keeping other parameter estimates constant, behaviour was simulated over the entire range of empirically found parameter values. The medians of the posterior distributions were just as point estimates for comparing to the generative parameter values. Crucially, a strong, statistically significant correlation between the generative and recovered volatility belief ($\omega$) parameters confirmed that our key model parameter could be recovered reliably ($r$=.92, 95% CI [0.87,0.93], $t(162)=40$, $p < 2.2\times10^{-16}$) (*Fig S3a*). Next, we tested recovery for our key belief state surprise ($\Im$) by calculating $\Im$ values from the inputs and the parameters and using them to simulate behaviour. We found strong positive correlations between the generative and recovered values for $\Im$ ($r$=.99, 95% CI[0.99, 0.99], $t(1971)=261$, $p < 2.2\text{x}10^{-16}$) (*Fig S3b*). Finally, we also confirmed the key response model response model beta-estimates could be recovered well; this was tested for post-error slowing (r=.96, 95% CI[ 0.96 , 0.97], $t(292)=66$, $p < 2.2\times10^{-16}$), expected uncertainty (r= .99, 95% CI[0.99, 0.99], r(292) = 319, $p < 2.2 \times 10^{-16}$), and unexpected uncertainty (r=.99, 95% CI[ 0.99, 0.99], t(321) = 2339, $p < 2.2 \times 10^{-16}$) (*Fig S3 c-e*).

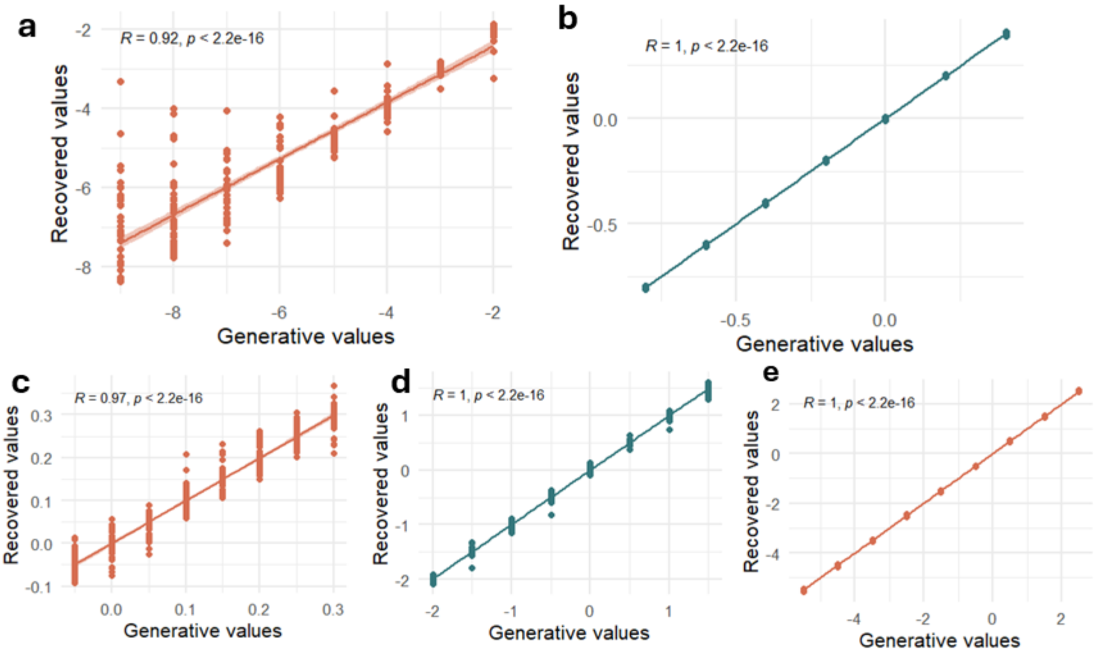


***Supplementary Figure S4.*** *Correlation between generative parameter values used to simulate the datasets (generative values) and the median posteriors recovered from the model-fitting (recovered values). a) Correlation between generative and recovered values for volatility beliefs parameter ($\omega$) (r=.92, p<2.2x $10^{-16}$) , b) Correlation between generative and recovered values for belief state surprise ($\Im$) (r=.99, p<2.2x $10^{-16}$), c) Correlation between generative and recovered values for response model beta estimate for post-error slowing ( r=.97, p<2.2x $10^{-16}$), d) Correlation between generative and recovered values for response model beta estimate expected uncertainty (r=.99, p<2.2x $10^{-16}$) , e) Correlation between generative and recovered values for response model beta estimate unexpected uncertainty(r=.99, p<2.2x $10^{-16}$).*
